# Temperature-dependent twist of double-stranded RNA probed by magnetic tweezers experiments and molecular dynamics simulations

**DOI:** 10.1101/2023.05.31.543084

**Authors:** Hana Dohnalová, Mona Seifert, Eva Matoušková, Flávia S. Papini, Jan Lipfert, David Dulin, Filip Lankaš

## Abstract

RNA plays critical roles in the transmission and regulation of genetic information and is increasingly used in biomedical and biotechnological applications. Functional RNAs contain extended double-stranded regions and the structure of double-stranded RNA (dsRNA) has been revealed at high-resolution. However, the dependence of the properties of the RNA double helix on environmental effects, notably temperature, is still poorly understood. Here, we use single-molecule magnetic tweezers measurements to determine the dependence of the dsRNA twist on temperature. We find that dsRNA unwinds with increasing temperature, even more than DNA, with Δ*Tw*_RNA_ = −14.4 ± 0.7 º/(°C·kbp), compared to Δ*Tw*_DNA_ = −11.0 ± 1.2 º/(°C·kbp). All-atom molecular dynamics (MD) simulations using a range of nucleic acid force fields, ion parameters, and water models correctly predict that dsRNA unwinds with rising temperature, but significantly underestimate the magnitude of the effect. These MD data, together with additional MD simulations involving DNA and DNA-RNA hybrid duplexes, reveal a linear correlation between twist temperature decrease and the helical rise, in line with DNA but at variance with RNA experimental data. We speculate that this discrepancy might be caused by some unknown bias in the RNA force fields tested, or by as yet undiscovered transient alternative structures in the RNA duplex. Our results provide a baseline to model more complex RNA assemblies and to test and develop new parameterizations for RNA simulations. They may also inspire physical models of temperature-dependent dsRNA structure.

## Introduction

Nucleic acid double helices in their DNA and RNA form play fundamental roles in biology. DNA is a carrier of genetic information in all cellular life. RNA performs critical functions in the transmission of genetic information and can adopt a variety of structures, with the double helix being a prominent structural motif [1]. Besides their biological role, DNA and RNA duplexes also serve as basic building blocks of artificial nanostructures [2, 3].

Life can flourish in a range of temperatures – extreme thermophiles can tolerate 100 °C, while extreme psychrophiles can survive at nearly 0 °C [4]. To thrive in a broad temperature range, organisms have to use efficient strategies of thermal adaptation. However, mechanisms of thermal adaptation at the molecular level in general, and thermal effects on nucleic acid properties and function in particular, are only starting to be understood [5-8]. Temperature-dependent properties of DNA and RNA double helices also play a role in nucleic acid nanostructures, which can operate in a broad temperature range and may even be thermally activated to perform functions related to biochemical diagnostics or drug delivery [9]. Thus, we need to better understand how the structure of nucleic acid double helices in their DNA and RNA form depends on temperature.

A number of studies have focused on temperature-dependent properties of double-stranded (ds) DNA. Thermal effects on dsDNA bending persistence length [10-12] and twist stiffness [13] have been examined using a variety of methods. Based on several lines of evidence, a two-state model of dsDNA structure and stiffness, including effects of temperature and other factors, has been formulated [14-16]. A pioneering study utilized atomic-resolution molecular dynamics (MD) simulations to probe temperature effects on dsDNA structure and elasticity [17]. In previous work, we have determined changes of dsDNA twist with temperature by magnetic tweezers (MT) measurements and quantitatively compared the experimental findings to atomic-resolution and coarse-grained MD simulations [18]. A follow-up MD study focused on temperature-dependent dsDNA bending and elongation [19]. A similar methodology combining MT measurements and MD simulations was used to examine the dsDNA twist dependence on the ionic environment [20]. The, at least in parts, quantitative agreement between MT measurements and all-atom MD data suggests that MD simulations of DNA oligomers ∼3 helical turns long, represented at atomic resolution and at the microsecond time scale, may offer a powerful and quantitative approach to probe thermal and ionic effects on DNA structure. While these works elucidated many aspects of temperature-dependent shape and stiffness of DNA duplexes, how structural properties of RNA double helices are affected by temperature remains largely unknown.

In this work, we examined the temperature dependence of twist of RNA and DNA double helices. We used magnetic tweezers (MT) to measure the temperature-dependent twist of dsRNA, and found that dsRNA twist decreases with temperature. The slope inferred from the experiment, −14.4 ± 0.7 º/(°C·kbp), is higher than the one for dsDNA, −11.0 ± 1.2 º/(°C·kbp), previously reported using the same experimental approach [18].

We complemented the experiments by atomic-resolution MD simulations of double-stranded RNA and DNA. To better understand the microscopic mechanism of twist temperature dependence, we also performed MD simulations of a DNA-RNA hybrid duplex. We simulated 33 base-pair (bp) oligomers, systematically testing several parameterizations of interatomic interactions (force fields) for the nucleic acid, water, and ions. For the dsDNA simulations, we employed the bsc1 force field [21] to extend our previous investigations [18] using the OL15 force field [22]. The RNA oligomer was simulated using four different force field combinations, while two force fields were used to simulate the hybrid.

Comparing the simulated data with experimental values, we found that both dsDNA force fields yield decreases of dsDNA twist with increasing temperature in quantitative agreement with the MT experiment. All the dsRNA simulations also indicate a decrease of dsRNA twist with temperature, agreeing qualitatively with the MT measurement. However, the simulated magnitude of the change dramatically deviates from the MT experiment – the dsRNA twist decrease inferred from MD is at least 3 times lower than the MT value. The MD data further suggest that the twist-temperature slopes of the double-stranded RNA, DNA or hybrid oligomer are tightly correlated with the oligomer compaction quantified by its helical rise. While this dependence is in line with experimental data for dsDNA, it again disagrees with dsRNA experimental values. We speculate that the discrepancy between experiment and simulation for the RNA duplex may be caused by some systematic bias in RNA force fields, or by the very different length and time scales probed in the MT experiment and in the MD simulations. The latter possibility would suggest the existence of as yet undisclosed transient structures within the RNA duplex which, contrary to known transient structures in dsDNA, would make the dsRNA twist thermal response length and time scale dependent. We discuss biological consequences of this possibility.

## Results

### Magnetic tweezers measurements reveal a strong decrease of double-stranded RNA twist with increasing temperature

We used a temperature-controlled high-throughput magnetic tweezers (MT) set-up [23] (**Materials and Methods**), where ∼3.3 kbp dsRNA molecules were tethered between the flow cell surface and 1 µm-diameter magnetic beads (**Fig. 1A**). Nick-free and end-labelled dsRNA constructs were generated by annealing of single-strands and subsequent ligation [24]. The dsRNA molecule is flanked by two handles, one randomly biotin-labelled at multiple points to attach to the magnetic bead and the other one randomly digoxygenin-labelled at multiple sites to bind to the anti-digoxygenin functionalized flow chamber glass surface. Permanent magnets mounted above the flow cell applied precisely calibrated stretching forces on the nucleic acid tether [25-29], by adjusting the height of the magnets above the flow cell, and to systematically over- and undertwist the tethered molecules, by rotating the magnets. We have previously shown that dsRNA exhibits an overall extension vs. applied rotation response very similar to dsDNA [30]. The dsRNA molecules were tested to determine their coilability preceding the experiment, as described in the **Materials and Methods**. We then measured extension vs. applied rotation curves at a low force of 0.3 pN, where the extension vs. rotation response is symmetric, i.e. the dsRNA molecule forms plectonemes for both negative and positive supercoiling (**Fig. 1AB**). We found that the extension vs. rotation curves systematically shift to lower turns with increasing temperature, from 0 turn at 25 °C to −3.4 turns at 50 °C (**Fig. 1B**). The systematic shift with temperature indicates that dsRNA unwinds when heated, a result qualitatively similar to DNA. Taking the mean value of the two independent analyses performed on the measurements (**Fig. 1CD**), we determined a change in dsRNA twist to be ∆*Tw*_*RNA*_ = (−14.4 0.7)°/(°C · kbp), i.e. dsRNA unwinds more than dsDNA since ∆*Tw*_*DNA*_ = (−11.0 ± 1.2)°/(°C · kbp) [18, 23]. Our value for ∆*Tw*_*RNA*_ is in excellent quantitative agreement with a recently published independent measurement [31], also using magnetic tweezers but a different dsRNA sequence, which reported ∆*Tw*_*RNA*_ = 15 ± 2 °/(°C · kbp) at the same ionic strength as our measurements and no dependence of the twist change with temperature on KCl concentration in the range 0.05 – 1 M, within experimental error.

**Figure 1.**
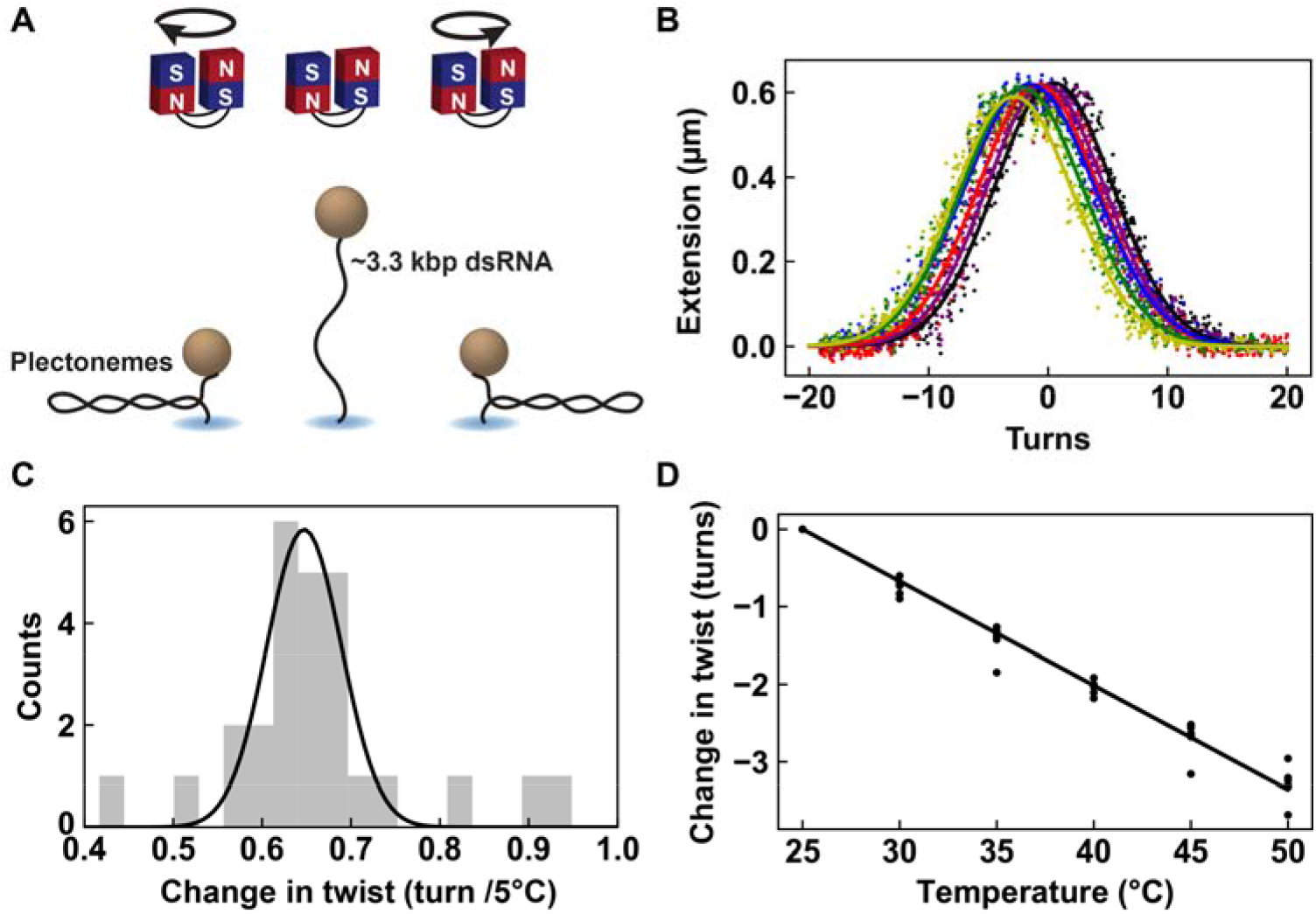
Single-molecule magnetic tweezers experiments reveal that dsRNA twist decreases with increasing temperature. **(A)** Schematic of the rotation-extension experiment performed with a magnetic tweezers instrument. The magnets are rotated from negative (left) to positive (right) turns, showing the formation of plectonemes upon addition of turns. **(B)** Rotation-extension traces performed at 25 °C (black), 30 °C (purple), 35 °C (blue), 40 °C (red), 45 °C (green) and 50 °C (yellow) for a single coilable dsRNA tether. The dots represent the 10-times decimated data and the solid lines are their respective Gaussian fits. **(C)** Distribution of the difference in turns at the maximum tether extension, i.e. the center of the rotation-extension curve, upon decreasing the temperature by 5 ºC, extracted from Gaussian fits to extension vs. rotation data for consecutive temperatures, for *N* = 6 independent dsRNA tethers. The solid line is a Gaussian fit with mean ± std of (0.65 ± 0.04) turns, corresponding to ∆*Tw*_*RNA*_ = (−14.1 + 1) °/(°C · kbp). **(D)** The position of the maximum tether extension as a function of temperature. The dots are measurements for *N* = 6 independent dsRNA tethers. The solid line is a linear fit with a slope that yields ∆*Tw*_*RNA*_ = (−14.6 + 1) °/(°C · kbp).

### Microsecond-scale MD simulations indicate a smaller decrease of dsRNA twist with temperature than the MT experiment

To understand the microscopic origin of the observed changes in twist of the dsRNA double helix with temperature and to quantitatively test available force fields for nucleic acids, we turned to all-atom molecular dynamics simulations. We performed four microsecond-long MD simulations of the same 33-bp dsRNA sequence, but employing different force fields for the nucleic acid, water and ions. In addition, we ran an analogous simulation of the dsDNA version of the same oligomer using the bsc1 force field for dsDNA, complementing the previously reported simulation [18] using the OL15 force field. The simulated systems are visualized in **Fig. 2AB** and the double-stranded DNA and RNA force field combinations employed (**Materials and Methods**) are listed in **Fig. 2** and **Table S1**.

**Figure 2.**
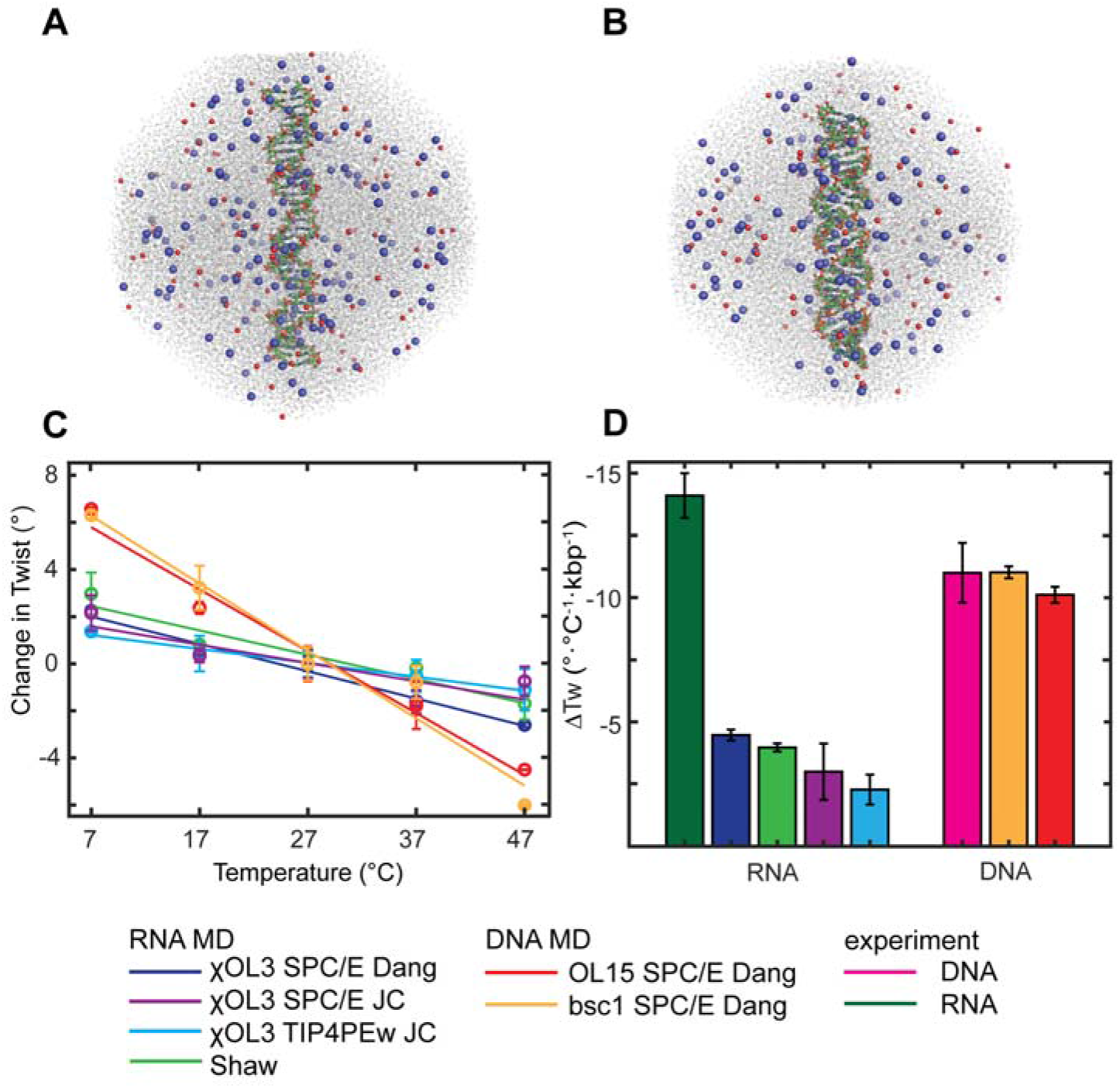
Molecular dynamics (MD) simulations of temperature-dependent double-stranded RNA and DNA twist compared to magnetic tweezers (MT) measurements. **(A**,**B)** The simulated systems containing 33 base-pair dsDNA **(A)** and dsRNA **(B)**, together with the K^+^ cations (blue), Cl^−^ anions (red) and water molecules (grey), are each immersed in an octahedral simulation box. The system sizes are shown approximately to scale – the dsRNA system is smaller, since the RNA duplex is shorter than the DNA one containing the same number of base pairs. **(C)** The simulated temperature changes of the end-to-end twist are well approximated by least-squares linear fits and all indicate a decrease of twist with rising temperature. **(D)** Comparison of the simulated twist-temperature slopes to magnetic tweezers measurements. While the MD data quantitatively agree with the dsDNA experiment, the MD simulations largely underestimate the measured twist decrease of the RNA duplex. The dsDNA MT experimental value is taken from [18].

Temperature changes of the end-to-end twist obtained from MD, together with the twist changes deduced from the MT experiments, are shown in **Fig. 2CD**, the numerical values with errors are in **Table S1**. The end-to-end twist for all the simulations decreases with temperature, the dependence being close to linear (**Fig. 2C**). The fitted MD temperature slopes and the experimental MT values are shown in **Fig. 2D**. The end-to-end twist decrease for dsDNA obtained using OL15 and bsc1 force fields are both within the error margins of the MT experiment. Thus, the quantitative agreement between the MT measurement and the OL15 simulations reported previously [18] is extended in this work also to the case of the bsc1 force field.

For dsRNA, all MD simulations again predict a decrease of the end-to-end twist with rising temperature, in qualitative agreement with the MT experiment. However, the magnitude of the simulated decrease is significantly underestimated compared to the MT experimental value. Indeed, the simulations indicate a change between -4.5 ± 0.2 and -2.3 ± 0.6 °/(°C.kbp) depending on the force filed, smaller than the dsDNA value and more than 3 times smaller than the experimental MT results (**Fig. 2CD, Table S1**). This large discrepancy common to all the force fields tested is further modulated by the differences between the individual force field parameterizations. The chiOL3 dsRNA force field, employed together with the SPC/E water model and the Dang ion parameters, yields the highest slope, closely followed by the Shaw force field with its recommended TIP4P-D water and CHARMM22 ions (**Fig. 2CD, Table S1**). The two simulations using the Joung-Cheatham ion parameters give a somewhat lower temperature slope, while the effect of the water model (three-point or four-point) is minor (**Fig. 2CD, Table S1**).

### Simulated twist temperature slopes tightly correlate with duplex compaction

To get further insight into the structural mechanism of the twist temperature change, we complemented our double-stranded RNA and DNA data by the MD simulations of a DNA-RNA hybrid. We again used a 33-bp oligomer of the same sequence as the DNA, but with T replaced by U in the complementary strand (**Materials and Methods**). Two force field combinations were tested: the RNA strand was modelled using the chiOL3 force field, while OL15 or bsc1 were used for the DNA strand. The inclusion of the hybrid oligomer enabled us to investigate a mechanism of twist temperature dependence common to all three simulated nucleic acid duplex variants.

**Fig. 3** shows the twist-temperature slope inferred from MD as a function of mean helical rise. It is seen that values for all the duplex variants (DNA, RNA and hybrid) and all the force fields examined follow a simple rule: the smaller the helical rise, the weaker the twist temperature decrease. Moreover, the relationship is very close to linear (*R*^2^ 0.98, straight line in **Fig. 3**). A smaller helical rise means shorter distance between base pairs measured along the helical axis, i.e. a shorter, more compact double helix. The MD simulations, therefore, suggest a mechanism of twist temperature dependence common to the DNA, RNA and hybrid duplexes: the more compact the duplex, the less sensitive its twist is to temperature changes.

**Figure 3.**
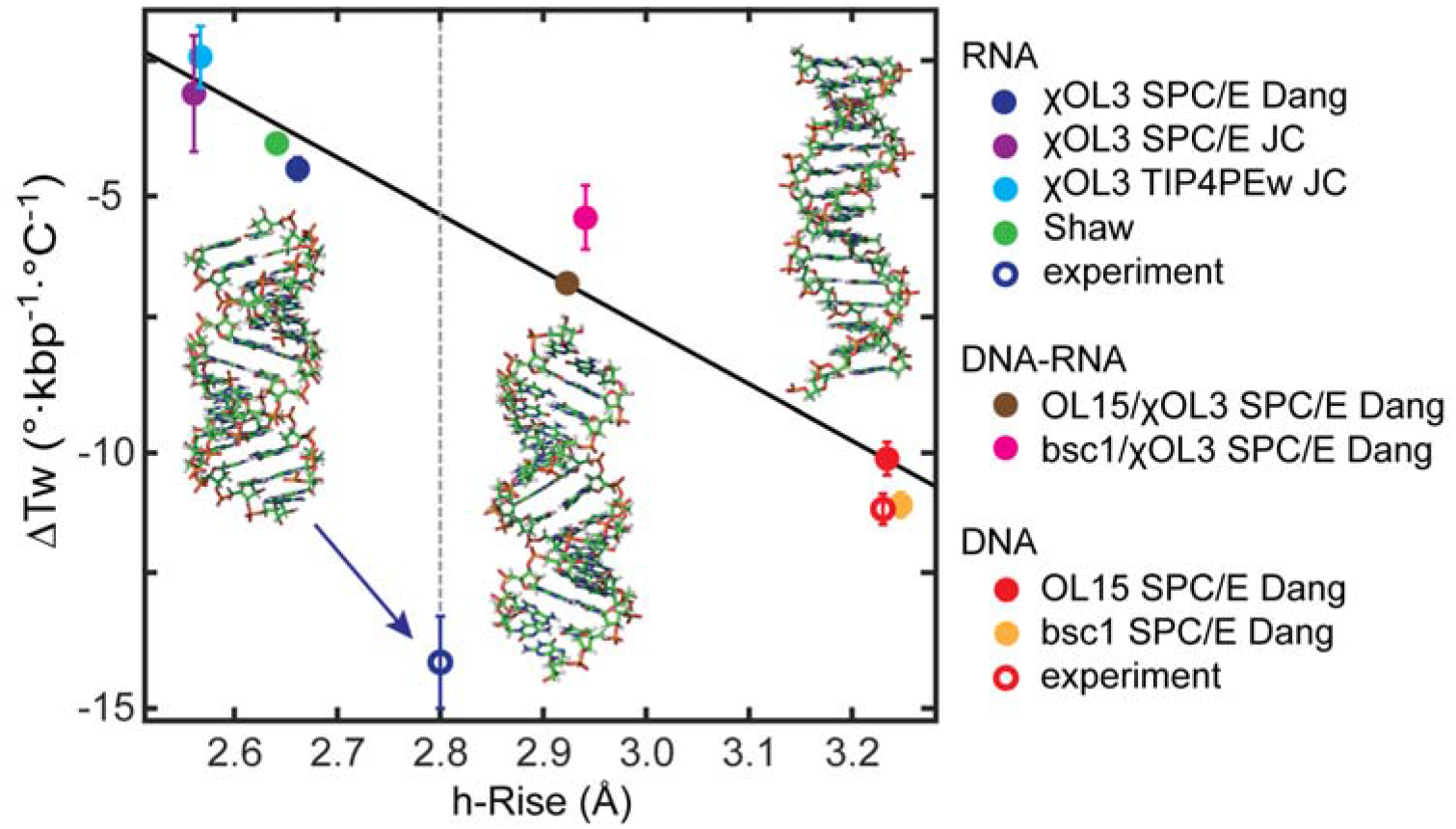
Temperature slopes of double-stranded DNA, RNA and hybrid DNA-RNA plotted against the mean helical rise. The dsDNA MT value from previous work [18] together with the h-rise from ASAXS measurements [32] (red empty circle), as well as the dsRNA MT value from this work and the consensus dsRNA h-rise [33] (empty blue circle highlighted by the blue arrow) are shown as experimental data points. The MD data shown as solid circles follow a linear relationship and agree with the DNA experimental data. However, the MD simulations disagree with the dsRNA experimental data that exhibit a much more negative twist temperature change, even stronger than for dsDNA. All the RNA MD simulations somewhat underestimate the helical rise, producing more compact structures than expected. But even if the h-rise is corrected to the consensus value (vertical broken line), the linear relationship would still yield the dsRNA twist-temperature slope much weaker than the MT experiment. Fragments of the 33-bp MD starting structures of dsDNA (right), the hybrid (middle) and dsRNA (left) are shown as well. Errors in MD h-rise values are very small and are omitted for clarity.

dsDNA, with the largest helical rise, also exhibits the largest twist decrease (**Fig. 3**, at the right). An experimental dsDNA data point (**Fig. 3**, red empty circle) is also shown. Its x-axis value is 3.23 Å, the midpoint of the distance between dsDNA base pairs in solution measured using anomalous small-angle X-ray scattering (ASAXS) with gold labels (3.23 ± 0.1 Å [32]), its y-axis value is the experimental MT result from [18]. The dsDNA experimental and simulated data are close to each other, demonstrating that microsecond-scale MD simulations can quantitatively reproduce not only the experimental twist-temperature slope but also the measured DNA helical rise in solution.

The situation is very different for dsRNA (**Fig. 3**, at the left). The experimental dsRNA data point in **Fig. 3** (blue empty circle, highlighted by an arrow) represents our MT value of -14.4 ± 0.7 °/(°C.kbp), together with the midpoint of the dsRNA consensus helical rise in solution containing monovalent ions (2.8 ± 0.1 Å, see [33] and references therein). All dsRNA MD simulations yield twist thermal change much smaller than the MT experiment, and they also somewhat underestimate the RNA helical rise compared to its solution consensus value. However, even if the underestimated MD helical rise is corrected to the consensus value, the linear relationship would imply the slope around -5 °/(°C.kbp), as indicated by the intersection between the fitting line and the 2.8 Å vertical line **(Fig. 3)**. This is still a much weaker effect than the measured value. As for the simulated hybrid DNA-RNA duplex, its helical rise is slightly lower than but close to the experimental value of ∼3.0 Å [34] and the twist temperature slope is around -6 °/(°C.kbp), both lying between the MD values for dsDNA and dsRNA.

Thus, while the linear model agrees quantitatively with experimental data for dsDNA, it is completely at odds with the dsRNA experimental data which indicate a much stronger twist temperature decrease, even if the model is corrected for the underestimated dsRNA helical rise.

### Effect of global twist definition

The MD results presented so far refer to the end-to-end twist as defined in the **Materials and Methods** – a twist angle between two right-handed, orthonormal frames located at the ends of the duplex. The twist angle was computed exactly as the local twist in the 3DNA algorithm [35], i.e. as the rotation angle between the end frames measured in the plane whose normal is the mean of the two z-axis vectors (mean plane). Taking just the end frames into consideration mimicked the MT experiment in the sense that it was the overall rotation between the two ends that mattered, while conformational features in the intervening part were not explicitly included. The 3DNA twist angle definition, moreover, ensured that the end-to-end twist changes were invariant with respect to constant offset rotations of the end frames about their z-axes, an offset that was also not known in the MT experiment [18].

To examine the dependence of our results on the global twist definition, we also determined the global twist as the sum of helical twists (h-twists) over all the base-pair steps involved. Helical twist definitions used by the 3DNA [35] and Curves+ [36] conformation analysis programs were tested. They are both based on the axis of the screw transformation mapping one base pair to the next one. However, while 3DNA uses the screw axis directly as the local helical axis, the Curves+ axis is smoothed using a polynomial weighting function. The temperature change of the sum of 3DNA h-twists (**Table S1**) underestimates the MT measurement already for dsDNA, as reported previously [18]. In the case of dsRNA, it is still further from the experimental value than the MD end-to-end twist, being even positive in one case (**Table S1**). The Curves+ data are closer to the MD end-to-end twist values (**Table S1**), in line with earlier results for dsDNA [18]. Nevertheless, the twist temperature slopes for the two h-twist definitions are both tightly correlated with the end-to-end twist data (**Fig. S1**). This is understandable – all three definitions ultimately depend on the relative rotations between bases and base pairs, and since the thermal effects are rather small, these dependencies can be linearized, yielding linear relations between the global twist values computed using any two of the definitions. Taken together, the various global twist definitions examined here consistently indicate a severe underestimation of dsRNA twist temperature change by MD simulations compared to the measured value.

## Discussion and Conclusions

In this work, we reported the changes of twist of nucleic acid double helices with temperature, probed by magnetic tweezers experiments and all-atom MD simulations. We first extended the previous work [18] to verify that both the current Amber force fields, bsc1 and OL15, yield dsDNA twist temperature change in quantitative agreement with the MT measurement. We then turned to the dsRNA and examined its twist temperature dependence by MT experiment and extensive MD simulations involving different force fields for dsRNA, water and ions. The duplex end-to-end twist inferred from the MD data decreased with temperature, agreeing qualitatively with the MT experiment. However, in contrast to the dsDNA case, we found a large discrepancy between the dsRNA twist temperature decrease measured by MT and its prediction by atomistic MD simulations, the latter being less than a third of the experimental value.

To obtain more insight into the microscopic origin of the twist temperature dependence, we complemented our MD data with simulations of a DNA-RNA hybrid oligomer, using two different force fields. The MD results including all three duplex variants (DNA, RNA, hybrid) and all the force fields revealed a tight linear correlation between the twist temperature slope and the duplex compaction quantified by the helical rise: more compact duplexes with smaller helical rise also exhibit lower sensitivity of twist to temperature changes. While this linear dependence agrees quantitatively with dsDNA experimental data, it predicts a much weaker twist thermal change for the dsRNA than experimentally observed, due to the much lower dsRNA twist temperature change deduced from MD.

The origin of such discrepancy is not *a priori* clear. One obvious possibility is some systematic shortcoming of the force fields tested. For instance, the (now obsolete) bsc0 Amber force field [37] examined in the prior study [18] underestimated the dsDNA twist temperature decrease by ∼32 %. Since the improvement to the current bsc1 and OL15 DNA force fields consists in adjusting the backbone torsional parameters, the possible dsRNA force field problem might be primarily related to the inaccurate description of the backbone. On the other hand, the discrepancy between MD and experiment observed here for dsRNA is much larger than in the case of the bsc0 DNA simulations. Furthermore, the deviation is similar for two entirely different force fields, chiOL3 and Shaw. Thus, it appears unlikely that the disagreement with the experiment is caused by a failure of a particular force field. If rooted in a force field bias at all, it might rather be due to general properties of this class of force fields, limited by their functional form and lack of polarization [38].

We note that a similar study using MT measurements and all-atom MD simulations to probe the dependence of dsDNA twist on both ion concentration and identity observed quantitative agreement for some, but also considerable deviations for other ions [20]. By contrast, the discrepancy between the simulated and experimental twist temperature change observed here is consistently large for all the ion and water models examined, adding confidence to the robustness of our results. The ability of MD force fields to faithfully reproduce structure, dynamics and elasticity of DNA and RNA duplexes has been extensively tested [38, 39]. For instance, modern dsDNA and dsRNA force fields are able to reproduce, both in sign and in magnitude, even such a subtle effect as the opposite coupling between twist and elongation in DNA and RNA duplexes, where DNA underwinds when stretched, while RNA overwinds [30, 40]. Furthermore, all the duplex variants (DNA, RNA and hybrid) and all the force fields tested here conform to the same linear relationship between the duplex helical rise and the twist temperature decrease (**Fig. 3**).

These considerations suggest a rather consistent picture, namely that the ∼30 bp double helices at the microsecond scale sample a certain domain of the conformational space, characterized by a tight correlation between the duplex helical rise and the sensitivity of its twist to temperature. In contrast to MD, the MT measurements take seconds to minutes and involve kilobase-long duplexes. Thus, they probe the double helix at much larger time and length scales than the MD simulations. There may be structural changes in the RNA duplex taking place at these longer scales, increasing the twist temperature dependence.

Such slow changes are well known to take place in the DNA duplex. They include the base-pair breathing at the 100 µs scale [41], formation of the Hoogsteen pairs at the millisecond scale [42], or the exchange between the *a* and *b* states in the – somewhat speculative – two-state model of DNA shape and stiffness [14-16]. Moreover, the changes may be cooperative, like in the two-state DNA model where the domain size exceeds 200 bp. Nevertheless, these processes do not seem to significantly affect the DNA twist change with temperature, whose measured value agrees quantitatively with the microsecond-scale MD prediction. By contrast, a large discrepancy is found in the case of the RNA double helix, still awaiting a possible structural explanation. We note that the MT measurements are performed at low stretching force (∼0.3 pN), that would presumably not preclude the formation of alternative structures.

Nevertheless, the microsecond time scale probed by MD does have relevance in the biological context. For instance, measured diffusion coefficients for transcription factors (TFs) range from ∼0.5 to 5 µm^2^/s, depending on their size, shape, and interactions with chromatin [43]. It follows that in a microsecond, a TF-like protein can travel a distance of 1.7 to 5.5 nm, comparable e.g. to the dsRNA diameter of 2.4 nm or its helical pitch of 3.2 nm. Thus, the properties at the microsecond scale may be relevant for the diffusion encounter, recognition and binding of these molecules to dsRNA, and our MD results suggest that at this scale, the RNA twist is only weakly affected by temperature. In contrast, the much larger twist temperature sensitivity at more extensive time and length scales as indicated by our MT measurements may affect properties of long stretches of dsRNA, such as in RNA viruses, where temperature is a critical factor defining the outcome of viral infections and the direction of viral evolution [8].

In summary, we have presented a combined experimental and simulation study to probe temperature-dependent twist of the RNA double helix. Both the magnetic tweezers measurements and all-atom MD simulations indicate that dsRNA unwinds with rising temperature. However, the magnitude of the twist temperature decrease observed in the simulations is much weaker than in the experiment. While some as yet undiscovered MD force field bias cannot be excluded, we also consider an alternative explanation, namely that this difference may reflect the existence of transient structures formed in the RNA duplex at time and length scales inaccessible to MD simulations. This would imply, in contrast to DNA, a scale-dependent structural response of the RNA double helix to temperature.

## Materials and Methods

### Magnetic tweezers measurement of dsRNA twist

The fabrication of the construct used in the magnetic tweezers experiments has been described in [24]. The temperature controlled high-throughput magnetic tweezers apparatus used in this study has been previously described in detail in [23]. Description of the flow cell assembly and preparation can be found in [24]. Specifically, 10 µl of Dynabeads MyOne Streptavidin T1 magnetic beads (Thermofisher, Germany, Cat # 65604D)) was washed twice in TE 1X buffer (10 mM Tris, 1mM EDTA pH 8.0, supplemented with 150 mM NaCl and 2 mM sodium azide), and subsequently mixed with ∼0.2 ng of coilable ∼3.3 kbp dsRNA and 1 mg/ml bovine serum albumin (BSA, New England Biolabs). The ∼3.3 kbp length includes only the stem, not the two ∼400 bp handles that attach the tether to the surface and the magnetic bead, respectively. The dsRNA attached beads were then washed once to remove the excess RNA, resuspended in 40 µl TE 1X buffer, and flushed in the flow cell. Following ∼10 minutes incubation, the excess magnetic beads were flushed out with 1 ml of TE 1X buffer, followed by flushing in of 1 ml phosphate saline buffer (PBS), in which the experiments took place. To determine whether the tethers were coilable, a test rotation-extension experiment was performed, rotating the magnets from -20 turns to +20 turns at 0.4 turns/s, and applying a force of 4 pN. A coilable molecule then shows no significant change in extension in negative supercoils, while its end-to-end extension decreases in positive supercoils [24, 30]. A non-coilable molecule shows no change in extension in both positive and negative supercoils, while a bead attached via multiple dsRNA tethers shows a decrease in extension upon applying both positive and negative turns.

To extract the dsRNA twist dependence on temperature, we performed dynamic rotation-extension experiments of the dsRNA in PBS at ∼0.3 pN force (**Fig. 1A**). At such low forces, the rotation-extension of a coilable dsRNA tether is symmetric, with an approximately Gaussian shape, where the maximum extension corresponds to the torsionally relaxed molecule [24, 30]. The data were recorded at 58 Hz camera acquisition frequency and the magnets were rotated at 0.4 turns/s from -20 to +20 turns. Dynamic rotation-extension experiments were repeated at each temperature, from 25 °C to 50 °C with incremental steps of 5 °C, as previously described for coilable dsDNA [23] (**Fig. 1B**). The rotation-extension traces where averaged 10-times, and subsequently fitted with a Gaussian function using a non-linear least-square fitting algorithm (Python 3) (**Fig. 1B**). The number of turns at maximum extension was extracted for each dsRNA tether from the Gaussian fit peak position, represented as a function of temperature with the value at 25 °C set to zero (**Fig. 1BC**), and fitted using a linear function using a least-square fitting algorithm (Python 3). From this fit, the dependence of the dsRNA twist changes on temperature was extracted (**Fig. 1D**).

### Molecular dynamics simulations and analysis

We simulated the 33-bp oligomer used in previous study [18], whose sequence of the reference strand reads GAGAT GCTAA CCCTG ATCGC TGATT CCTTG GAC. The RNA version of the duplex has a sequence where U replaces T in both strands, the hybrid sequence was obtained by replacing T by U in the complementary strand only. We performed atomic-resolution MD simulations with explicitly represented water molecules and ions, using the Amber17 suite of programs. Each system was simulated at 7, 17, 27, 37 and 47 °C. A new DNA simulation using the bsc1 force field [21] was produced, complementing the OL15 [22] simulation reported previously [18], while the chiOL3 [44, 45] and Shaw [46] force fields were used for RNA. The Dang [47] and Joung-Cheatham (JC) [48] ion parameters, and the SPC/E as well as TIP4PEw water models were utilized, with the exception of the Shaw RNA force field which was combined with its recommended CHARMM22 ion parameters [49] and the TIP4P-D water model [50]. The parameter combinations utilized are shown in **Fig. 2** and **Table S1**. The TIP3P water model was excluded due to its limited capability to reproduce properties of real water and their temperature dependence [51].

The DNA and RNA duplexes were built in their canonical B- and A-form, respectively, using the *nab* module of Amber. To build the hybrid, the 3D-NuS server [52] with the *sequence specific model* and *nmr* options was used. The systems were immersed in an octahedral periodic box containing the duplex, water molecules, K+ ions to neutralize the duplex charge, and additional K+ and Cl-ions to mimic the physiological concentration of 150 mM KCl (**Fig. 2AB**). They were then subjected to a series of energy minimizations and short MD runs before starting the production of MD trajectories, 1µs each. The standard Amber hydrogen mass repartitioning and the time step of 4 fs were employed. Snapshots were taken every 10 ps. The oligomer global twist was measured as the end-to-end, or mean-plane twist between reference frames at the oligomer ends and was defined exactly as the local twist between neighbouring base pairs in the 3DNA algorithm [35]. The end frames were obtained by projecting the end base-pair frames (defined as in 3DNA) onto the local helical axis, computed as the mean axis of the two steps containing the pair. The helical axes of the steps were again calculated as in 3DNA. The changes of the end-to-end twist defined in this way are invariant with respect to the constant offset rotation of the end frames around their z-axes [18]. We also probed the global twist defined as the sum of helical twists of the base-pair steps within the analysed fragment, either computed as in 3DNA or extracted from the output of the Curves+ conformational analysis program [36]. Details of the protocol can be found in the previous work [18]. Only the inner 27 bp were analysed, 3 bp at each end were excluded. The errors were estimated as the mean absolute difference between the value for the whole trajectory and for its halves.

## Acknowledgement

We thank Willem Vanderlinden and Nadine Schwierz for useful discussions, and Marie Zgarbová for her help in preparing the simulations. DD was supported by the Interdisciplinary Center for Clinical Research (IZKF) at the University Hospital of the University of Erlangen-Nuremberg, the German Research Foundation grant DFG-DU-1872/3-1, DFG-DU-1872/4-1, DFG-DU-1872/5-1 and BaSyC – Building a Synthetic Cell” Gravitation grant (024.003.019) of the Netherlands Ministry of Education, Culture and Science (OCW) and the Netherlands Organisation for Scientific Research (NWO). HD and FL were supported by the grant of Specific University Research provided by the University of Chemistry and Technology Prague (grant no. A2_FCHT_2020_047).

## Notes

### Competing Interest Statement

The authors have declared no competing interest.

