## Supplemental Information for "Temperature-dependent twist of double-stranded RNA probed by magnetic tweezers experiments and molecular dynamics simulations"

**Table S1. Change of twist with temperature obtained from magnetic tweezer (MT) experiments and molecular dynamics (MD) simulations.**

| | | $\Delta Tw/\Delta T^a$ ($deg \cdot °C^{-1} \cdot kbp^{-1}$) | $\Delta hTw/\Delta T^b$ ($deg \cdot °C^{-1} \cdot kbp^{-1}$) | $\Delta hTw/\Delta T^c$ ($deg \cdot °C^{-1} \cdot kbp^{-1}$) |
|---|---|---|---|---|
| **RNA** | experiment | -14.4 ± 0.7 | | |
| | χol3 SPC/E Dang | -4.5 ± 0.2 | -4.1 ± 0.1 | -1.5 ± 0.2 |
| | Shaw | -4.0 ± 0.2 | -3.7 ± 0.1 | -0.8 ± 0.1 |
| | χol3 SPC/E JC | -3.0 ± 1.1 | -2.0 ± 1.1 | -0.1 ± 1.2 |
| | χol3 TIP4PEw Dang | -2.3 ± 0.6 | -2.0 ± 0.4 | -0.7 ± 0.3 |
| **DNA** | experiment[d] | -11.0 ± 1.2 | | |
| | bsc1 SPC/E Dang | -11.0 ± 0.2 | -11.3 ± 0.4 | -7.9 ± 0.4 |
| | OL15 SPC/E Dang | -10.1 ± 0.3 | -9.8 ± 0.3 | -6.9 ± 0.4 |
| **DNA-RNA** | bsc1/ χol3 SPC/E Dang | -5.4 ± 0.6 | -4.7 ± 0.9 | -2.0 ± 0.4 |
| | OL15/ χol3 SPC/E Dang | -6.7 ± 0.1 | -6.9 ± 0.2 | -3.7 ± 0.4 |

[a] MD values obtained using the end-to-end twist. Since this global twist definition mimics the MT setup, the experimental values are reported in the same column.

[b] MD values obtained using the sum of Curves+ helical twists.

[c] MD values obtained using the sum of 3DNA helical twists.

[d] Experimental value from ref. [1].

[1] Kriegel F, Matek C, Drsata T, Kulenkampff K, Tschirpke S, Zacharias M, et al. The temperature dependence of the helical twist of DNA. Nucleic Acids Res. 2018;46:7998-8009.

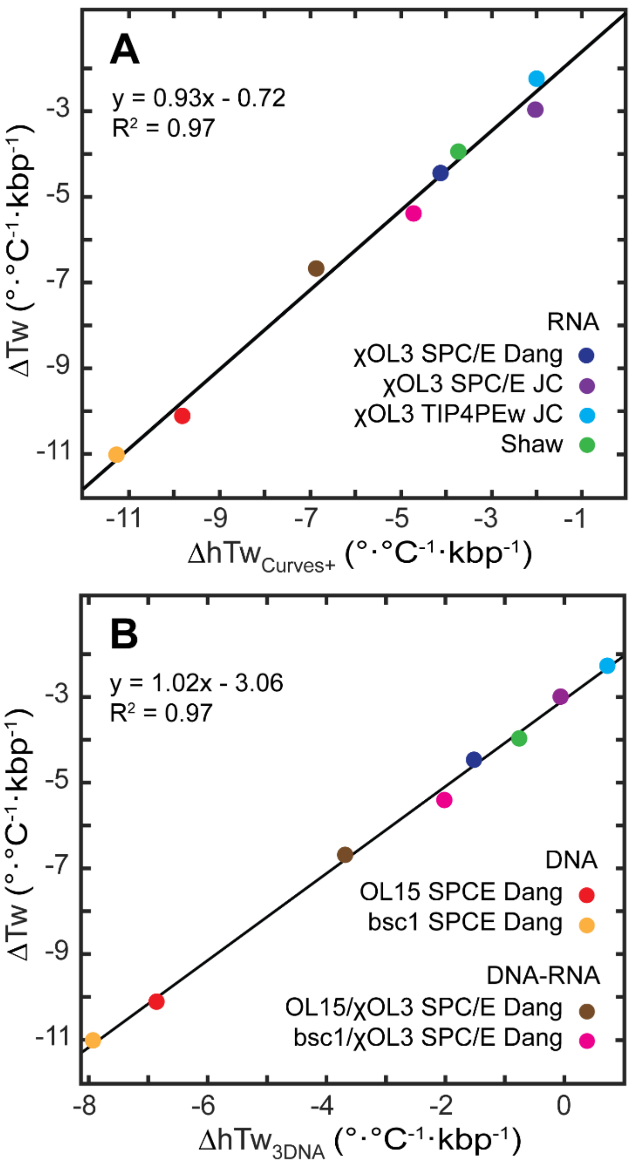


**Figure S1. Relations between twist-temperature slopes computed using various definitions of the oligomer global twist. (A)** End-to-end twist and sum of helical twists (h-twists) extracted from the Curves+ output, **(B)** end-to-end twist and sum of h-twists computed as in 3DNA. The equation of the fitting line and the square correlation coefficient are also shown. The colour coding indicated holds for both panels.
